# Combining multiple stressors unexpectedly blocks bacterial migration and growth

**DOI:** 10.1101/2024.05.27.595753

**Authors:** Anuradha Sharma, Alexander M. Shuppara, Gilberto C. Padron, Joseph E. Sanfilippo

**Affiliations:** Department of Biochemistry, University of Illinois at Urbana-Champaign, Urbana, IL, USA, 61801

## Abstract

In nature, organisms experience combinations of stressors. However, laboratory studies typically simplify reality and focus on the effects of an individual stressor. Here, we use a microfluidic approach to simultaneously provide a physical stressor (shear flow) and a chemical stressor (H_2_O_2_) to the human pathogen *Pseudomonas aeruginosa*. By treating cells with levels of flow and H_2_O_2_ that commonly co-occur in nature, we discover that previous reports significantly overestimate the H_2_O_2_ levels required to block bacterial growth. Specifically, we establish that flow increases H_2_O_2_ effectiveness 50-fold, explaining why previous studies lacking flow required much higher concentrations. Using natural H_2_O_2_ levels, we identify the core H_2_O_2_ regulon, characterize OxyR-mediated dynamic regulation, and dissect the redundant roles of multiple H_2_O_2_ scavenging systems. By examining single-cell behavior, we serendipitously discover that the combined effects of H_2_O_2_ and flow block pilus-driven surface migration. Thus, our results counter previous studies and reveal that natural levels of H_2_O_2_ and flow synergize to restrict bacterial colonization and survival. By studying two stressors at once, our research highlights the limitations of oversimplifying nature and demonstrates that physical and chemical stress can combine to yield unpredictable effects.

## Introduction

Traditionally, bacterial stress responses have been studied in reductionist laboratory conditions that poorly replicate complex host environments. Decades of work has elucidated the mechanisms that underly bacterial responses to isolated stressors such as oxidative stress (1, 2), nutrient deprivation (3), pH (4), and temperature (5). However, there remains a great challenge in understanding how these responses interrelate to defend against unique combinations of stressors. Furthermore, combining stressors in a test tube fails to capture the single-cell dynamics of stress responses found in natural environments. Recent studies from our lab and others have used microfluidic technology to simultaneously trigger and measure bacterial stress responses with single-cell resolution (6–12).

Fluid flow is central to the function of many host systems (13–15). In recent years, bacterial responses to flow have been explored using microfluidic devices. While there is evidence that bacteria respond to shear forces associated with flow (16–18), there is also a growing body of literature that shows that bacteria respond to flow-driven chemical transport (6, 19, 20). Logically, it is easy to understand how flow physically impacts local chemical environments by washing away or replenishing small molecules. However, the biological consequences of flow remain unclear and largely untested. So far, flow has been shown to have contrasting biological effects, from blocking quorum sensing (19) to promoting chemical stress (6). Based on these results, it is difficult to predict how bacteria will respond to a particular flowing environment. As flow is a defining feature of many host systems, there is critical need to study bacterial behavior in host-relevant flow.

H_2_O_2_ is found at micromolar levels in host tissues (21–23). Bacterial responses to H_2_O_2_ have been extensively studied in traditional lab conditions (24–30). Bacteria defend themselves against the harmful effects of H_2_O_2_ with catalases (encoded by *kat* genes) that enzymatically break down H_2_O_2_ (26). Bacteria also protect against H_2_O_2_ with NADH peroxidases (encoded by *ahp* genes) that use the reducing power of NADH to convert H_2_O_2_ into water (31). Many bacterial species directly sense intracellular H_2_O_2_ using OxyR, a transcription factor that directly activates expression of *kat* and *ahp* genes (26, 27, 32, 33). Although the mechanisms underlying H_2_O_2_ stress responses are largely known, almost all research has used H_2_O_2_ concentrations orders of magnitude higher than those found in host environments (34–37). Furthermore, research on many diverse bacterial species has established that millimolar concentrations (∼1,000 fold higher than found in hosts) are required to inhibit bacterial growth (38–42). Thus, the impact of micromolar H_2_O_2_ on bacteria is unclear and there is a critical need to investigate H_2_O_2_ responses in conditions that more precisely model the host environment.

What happens when bacteria encounter two simultaneous stressors? While it is intuitive to understand that two stressors are likely worse than one stressor, the interaction effects are often unpredictable. In a simple case, two stressors could have an additive effect, where the combination is twice as bad as either stressor alone. In a more complex case, two stressors could exhibit positive synergy, where the combination is more than twice as bad as either stressor alone. Here, we use microfluidics and single-cell imaging to explore the interactive effects of host-relevant flow and H_2_O_2_ on the human pathogen *P. aeruginosa*. Our results reveal that flow and H_2_O_2_ act synergistically to inhibit surface migration and growth.

## Results

To understand how multiple host stressors combine to restrict bacterial pathogens, we first aimed to establish how host-relevant H_2_O_2_ impacts the human pathogen *P. aeruginosa*. While human hosts contain micromolar concentrations of H_2_O_2_ (21–23), almost all experiments studying the effects of H_2_O_2_ on *P. aeruginosa* have used millimolar concentrations (35–37). Thus, we became interested in testing the effects of low micromolar doses on *P. aeruginosa* physiology. To test the impact of low H_2_O_2_ doses, we performed RNA-sequencing on *P. aeruginosa* cells after exposure to 8 µM H_2_O_2_ (Figure 1). For this experiment, we collected RNA after 5 minutes of H_2_O_2_ treatment, which allowed us to identify the most direct targets of H_2_O_2_-sensitive gene regulation. Our experiment revealed 17 upregulated genes and 3 downregulated genes after H_2_O_2_ treatment (Figure 1A). Our results demonstrate that host-relevant micromolar H_2_O_2_ doses are sufficient to alter gene expression of *P. aeruginosa*.

**Figure 1:**
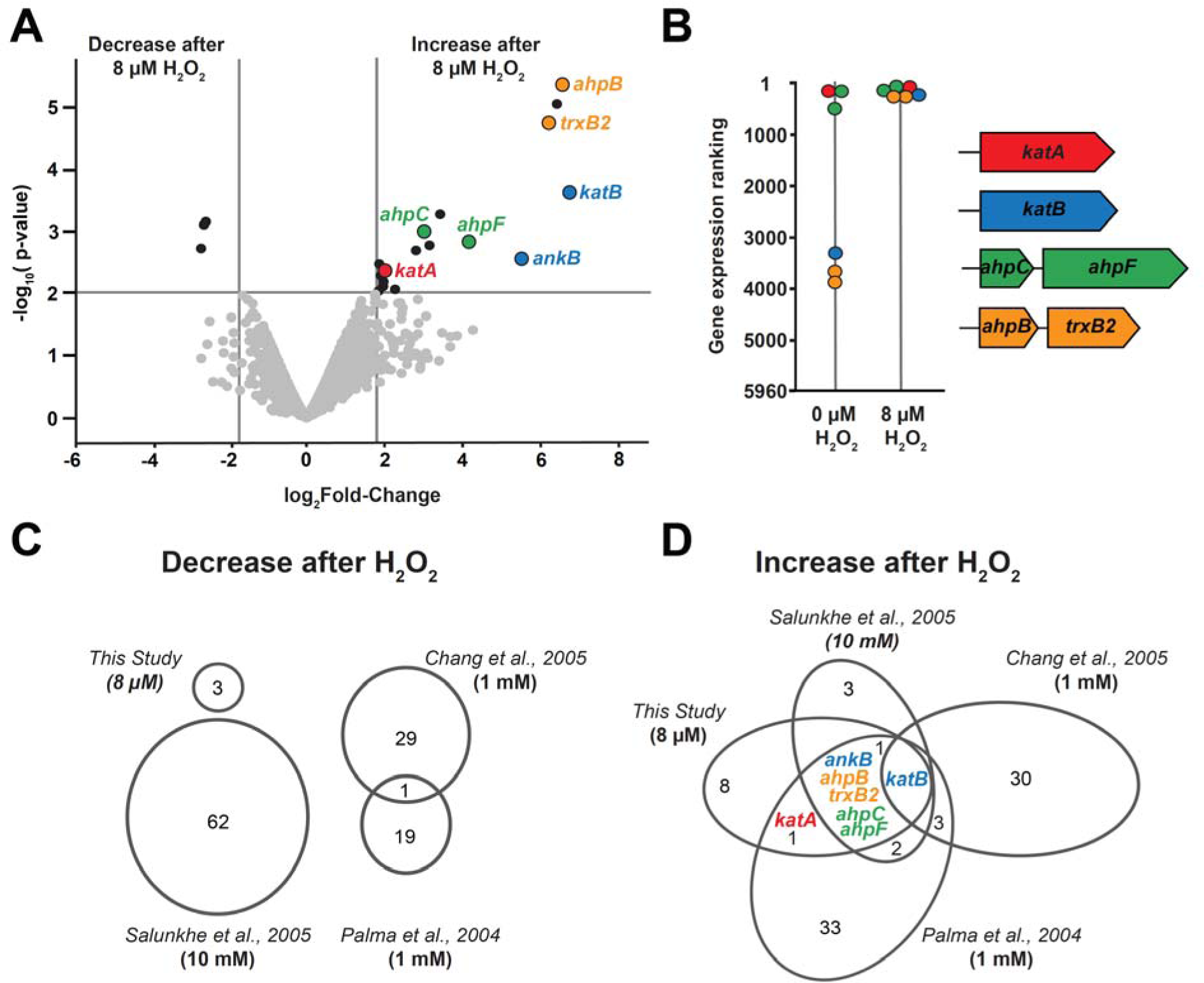
*P. aeruginosa* induces a small, clearly-defined H_2_O_2_ regulon. **(A)** RNA Sequencing shows that 17 genes are up-regulated while 3 genes are down-regulated in *P. aeruginosa* cells exposed to 8 µM H_2_O_2_ for 5 minutes. **(B)** Gene rankings based on relative transcript abundance with and without 8 µM H_2_O_2_ treatment. H_2_O_2_-sensitive genes shift to the top 150 highly expressed genes after H_2_O_2_ exposure. Gene organization of *katA*, *katB, ahpCF*, and *ahpB-trxB2*. (**C)** Comparative analysis of down-regulated genes from our study and three other transcriptomic studies. P-value (calculated using a hypergeometric distribution) for the 1 gene overlap between 2 studies is >0.05, demonstrating that the overlap is not significant. **(D)** Comparative analysis of up-regulated genes from our study and three other transcriptomic studies. Genes denoted with colors are known or predicted to have roles in H_2_O_2_ protection.

To explore if our experiment identified the same genes as previous studies, we compared our results to the available data from three independent microarray experiments (35–37) that tested *P. aeruginosa* gene expression in response to H_2_O_2_. All three previous studies used 1 or 10 mM H_2_O_2_ concentrations, which are approximately 1,000-fold higher than our experiment. As genes that are truly downregulated by H_2_O_2_ should appear in multiple studies, we were surprised to find no downregulated genes shared by all 4 datasets (Figure 1C). Additionally, no downregulated genes were shared by 3 datasets and only 1 downregulated gene was shared by 2 datasets (Figure 1C). Furthermore, a statistical analysis revealed that the 1 gene shared between 2 datasets was due to chance, suggesting that no genes are truly downregulated in response to H_2_O_2_.

How many genes are truly upregulated by *P. aeruginosa* in response to H_2_O_2_? Reasoning that genes truly upregulated by H_2_O_2_ should appear in multiple studies, we examined how the results of our study and the three previous H_2_O_2_ microarray studies overlap (35–37). While 110 genes were reported as upregulated in at least one study, our comparative analysis revealed only 14 upregulated genes that appeared in multiple studies (Figure 1D). Based on these results, we hypothesized that the three microarray datasets overrepresented the number of genes regulated by H_2_O_2_ due to their use of very high H_2_O_2_ doses. Supporting our hypothesis, our use of host-relevant micromolar H_2_O_2_ identified most (9 of 14) of the overlapping genes (Figure 1D). Thus, in contrast to all previous studies (35–37), we conclude that *P. aeruginosa* induces a small H_2_O_2_-sensitive regulon.

To characterize the *P. aeruginosa* H_2_O_2_ regulon, we identified genes that met two criteria: induction by host-relevant H_2_O_2_ (Figure 1A) and representation in multiple studies (Figure 1D). Of the 9 genes that met these two criteria, 7 have been implicated in H_2_O_2_ protection. To understand how the expression levels of these 7 genes compared to the rest of genome, we ranked all 5,960 *P. aeruginosa* PA14 genes based on relative transcript abundance (Figure 1B). Before H_2_O_2_ treatment, 2 genes (*katA* and *ahpC*) are among the 100 most highly expressed genes (Figure 1B). Both of these genes are located close to the origin of replication (ori), suggesting that they have critical roles in H_2_O_2_ protection (Figure S1). The other 5 genes (*ahpF*, *katB*, *ankB*, *ahpB*, *trxB2*) are among the 150 most highly expressed genes following 8 µM H_2_O_2_ treatment, suggesting that they also have important roles in H_2_O_2_ protection (Figure 1B)(Table S4). Together, our results indicate that *P. aeruginosa* induces a small, highly-expressed suite of genes in response to host-relevant H_2_O_2_ stress.

Many of the proteins upregulated by *P. aeruginosa* are known to protect cells against the harmful effects of H_2_O_2_. KatA and KatB function as catalases, which break down H_2_O_2_ into water and O_2_ (Figure S1) (26). AhpC and AhpF works as a pair to reduce H_2_O_2_ to water using NADH as a reductant (Figure S1) (26, 31). In *P. aeruginosa* and other bacteria, H_2_O_2_ sensing is mediated by the H_2_O_2_-sensitive transcription factor OxyR (26, 27, 32). To examine the role of OxyR in regulating our 7 genes, we scanned the relevant promoter regions for OxyR binding sites. Our 7 genes cluster into 4 distinct genomic locations (Figure S1), indicating the presence of at least 4 H_2_O_2_-sensitive promoters. Our search revealed OxyR binding sites in all 4 promoter regions (Figure S2), suggesting that OxyR plays a central role regulating the *P. aeruginosa* H_2_O_2_ regulon. To experimentally test the role of OxyR, we measured RNA levels of *katB* (the only gene upregulated in all four studies) and confirmed that *katB* levels are regulated by H_2_O_2_ and OxyR (Figure S3). Collectively, our data support a model where OxyR directly senses H_2_O_2_ and transcriptionally induces a small suite of proteins to protect *P. aeruginosa* against host-relevant H_2_O_2_ stress.

How does H_2_O_2_ combine with other host-relevant stressors to impact bacteria? During host colonization, bacteria typically experience environments where H_2_O_2_ and shear flow co-occur. To examine the impact of H_2_O_2_ and shear flow, we used microfluidics to simultaneously treat cells with shear flow (800 sec^-1^) and host-relevant H_2_O_2_ (8 µM). We introduced P*_katB_*::YFP reporter cells into our microfluidic devices to examine *katB* transcriptional activity. Our reporter also contained a constitutively expressed mCherry, which we used for normalization during the quantification process. In response to 8 µM H_2_O_2_ and 800 sec^-1^ flow, we observed that *katB* levels are dynamically induced, peaking at 40 minutes before returning to basal levels (Figure 2B, 2C). To test if *katB* dynamics are controlled by OxyR, we measured P*_katB_*::YFP reporter activity in an Δ*oxyR* mutant and observed no induction (Figure 2C). Thus, our results indicate that *P. aeruginosa* cells use OxyR to regulate a dynamic transcriptional response to the combined effects of H_2_O_2_ and flow.

**Figure 2:**
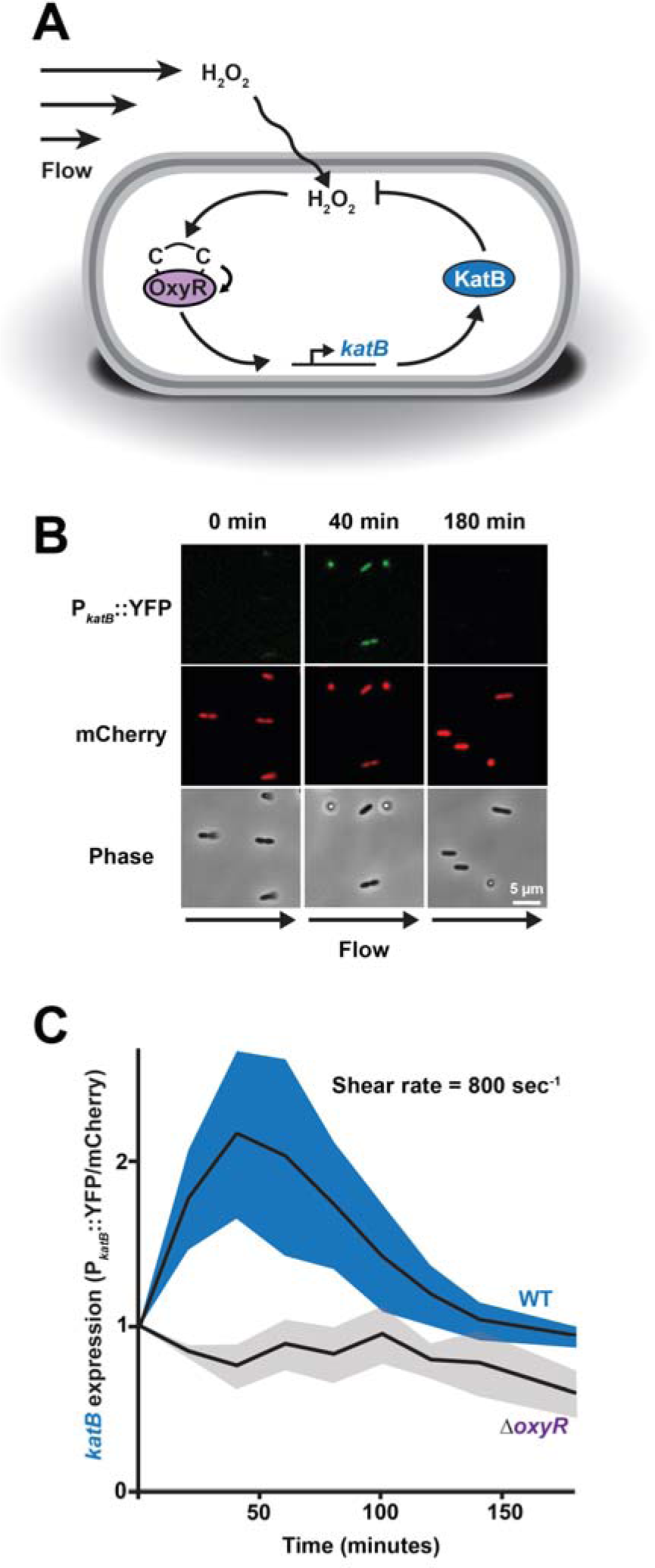
Host-relevant H_2_O_2_ triggers dynamic OxyR-mediated gene expression. **(A)** Model of OxyR-mediated *katB* expression in response to shear flow and H_2_O_2_. OxyR directly senses incoming H_2_O_2_ through reactive cysteine residues and activates transcription of *katB*. KatB functions as a catalase that degrades H_2_O_2,_ forming a negative feedback loop. **(B)** P*_katB_*:: YFP fluorescence, mCherry fluorescence (for normalization), and phase images of the *P. aeruginosa katB* reporter strain exposed to 8 µM H_2_O_2_ in flow at a shear rate of 800 sec^−1^ over 180 minutes. Scale bar is 5 μm. **(C)** Quantification of *katB* transcriptional reporter in response to 8 µM H_2_O_2_ over time for wild-type cells (blue) and Δ*oxyR* mutant cells (gray). *katB* transcriptional activity was quantified as YFP divided by mCherry. Shaded regions show SD of three biological replicates.

While investigating *katB* dynamics in flow, we noticed that *P. aeruginosa* cells exposed to H_2_O_2_ did not exhibit normal surface migration. *P. aeruginosa* cells use extension and retraction of type IV pili to migrate along solid surfaces through a process known as twitching motility. In flow, twitching motility proceeds against flow due to the passive reorientation of cells such that their type IV pili face upstream (43). To test the impact of H_2_O_2_ on surface migration in flow, we treated *P. aeruginosa* cells with a range of H_2_O_2_ doses in microfluidic devices. We simultaneously examined individual cells and quantified the percentage of cells that were twitching (Figure 3). While most (>80%) untreated cells twitched, increasing H_2_O_2_ concentrations led to an exponential decrease in twitching (Figure 3A). H_2_O_2_ concentrations of 16 µM or greater inhibited most cells (>80%) from twitching (Figure 3A, 3B). Our results demonstrate that host-relevant H_2_O_2_ blocks twitching motility of *P. aeruginosa* in flow.

**Figure 3:**
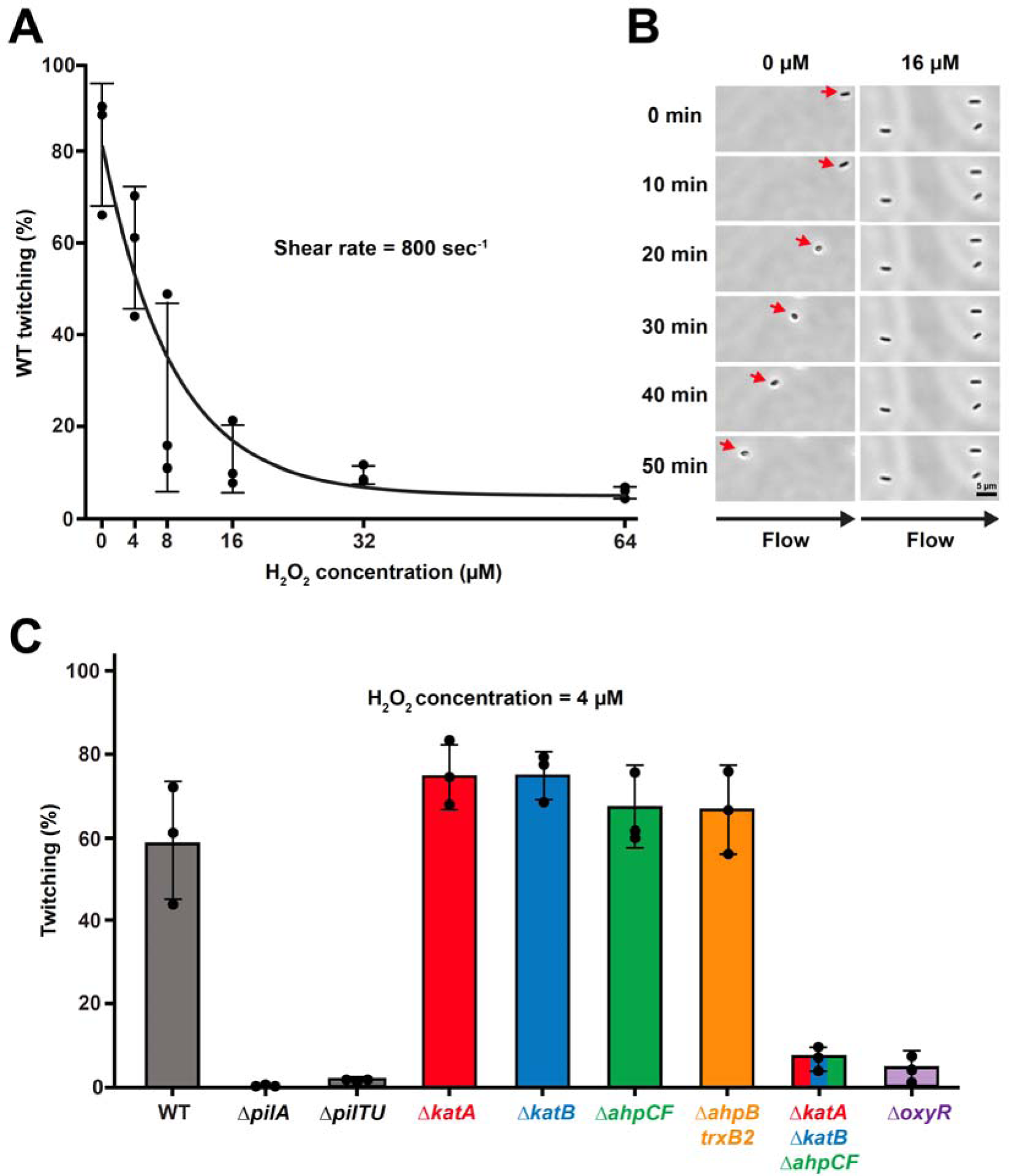
Host-relevant H_2_O_2_ doses inhibit *P. aeruginosa* surface migration in flow. **(A)** Quantification of the relationship between H_2_O_2_ concentration (0-64 µM) and twitching motility of *P. aeruginosa*. **(B)** Images representing twitching motility of *P. aeruginosa* over 50 minutes with no H_2_O_2_ and 16 µM H_2_O_2_. Without H_2_O_2_, wild-type cells twitch upstream whereas with 16 µM H_2_O_2_, wild-type cells do not twitch. Scale bar is 5 μm. **(C)** Quantification of twitching in response to 4 µM H_2_O_2_. Wild-type cells (gray) twitch, while Δ*pilA* and Δ*pilTU* cells (gray) serve as negative controls and do not twitch. Δ*katA* (red), Δ*katB* (blue), Δ*ahpCF* (green), or Δ*ahpB*-*trxB2* (yellow) mutant cells exhibit twitching motility indistinguishable from wild-type cells. Δ*katA* Δ*katB* Δ*ahpCF* (red, blue, and green) and Δ*oxyR* (purple) mutant cells are impaired at twitching. All twitching experiments were performed in flow at a shear rate of 800 sec^-1^. For each biological replicate, at least 45 cells were observed for quantification. Error bars represent SD of three biological replicates.

Is the H_2_O_2_ stress response required for *P. aeruginosa* twitching motility in flow? To test the impact of H_2_O_2_-sensitive genes, we quantified twitching of *P. aeruginosa* cells in a microfluidic device treated with 4 µM H_2_O_2_ and 800 sec^-1^ flow (Figure 3C). While 60 percent of wild-type cells were capable of twitching, Δ*pilA* mutant cells (which lack pili) and Δ*pilTU* mutant cells (which lack pilus retraction) exhibited no twitching and served as negative controls (Figure 3C). In contrast, mutants lacking *katA*, *katB*, *ahpCF,* or *ahpB*-*trxB2* exhibited twitching motility indistinguishable from wild-type (Figure 3C), leading us to hypothesize that *P. aeruginosa* uses redundant systems to protect against H_2_O_2_. In support of this hypothesis, almost no Δ*katA* Δ*katB* Δ*ahpCF* mutant cells (<10%) exhibited twitching (Figure 3C). Also, we observed that almost no Δ*oxyR* mutant cells (<10%) exhibited twitching (Figure 3C), indicating that the ability to respond to H_2_O_2_ is required for *P. aeruginosa* to migrate on surfaces in flow.

In light of our discovery that H_2_O_2_ blocks twitching, we wondered if host-relevant H_2_O_2_ has a general detrimental impact on *P. aeruginosa* physiology in flow. To investigate the impact of H_2_O_2_ and flow on cell growth, we examined *P. aeruginosa* cells growing in microfluidic devices exposed to a wide range of H_2_O_2_ concentrations (Figure 4). During infection of the lungs, bloodstream, or urinary tract, *P. aeruginosa* cells experience H_2_O_2_ concentrations of approximately 1-25 µM (21–23). While *P. aeruginosa* cells exhibited robust growth up to 100 µM H_2_O_2_ without flow, concentrations as low as 8 µM inhibited growth in flow (Figure 4A,S4). To dissect the flow dependency of growth inhibition, we measured single-cell growth in microfluidic devices with 8 µM H_2_O_2_ and shear rates of 80 – 800 sec^-1^, which are found in the human bloodstream (Figure 4B) (13). Low flow (80 sec^-1^) allowed for robust growth, medium flow (240 sec^-1^) partially inhibited growth, and high flow (800 sec^-1^) completely inhibited growth (Figure 4B, 4C). These results demonstrate that the combinatorial effects of H_2_O_2_ and shear flow are sufficient to block *P. aeruginosa* growth in host-relevant environments.

**Figure 4:**
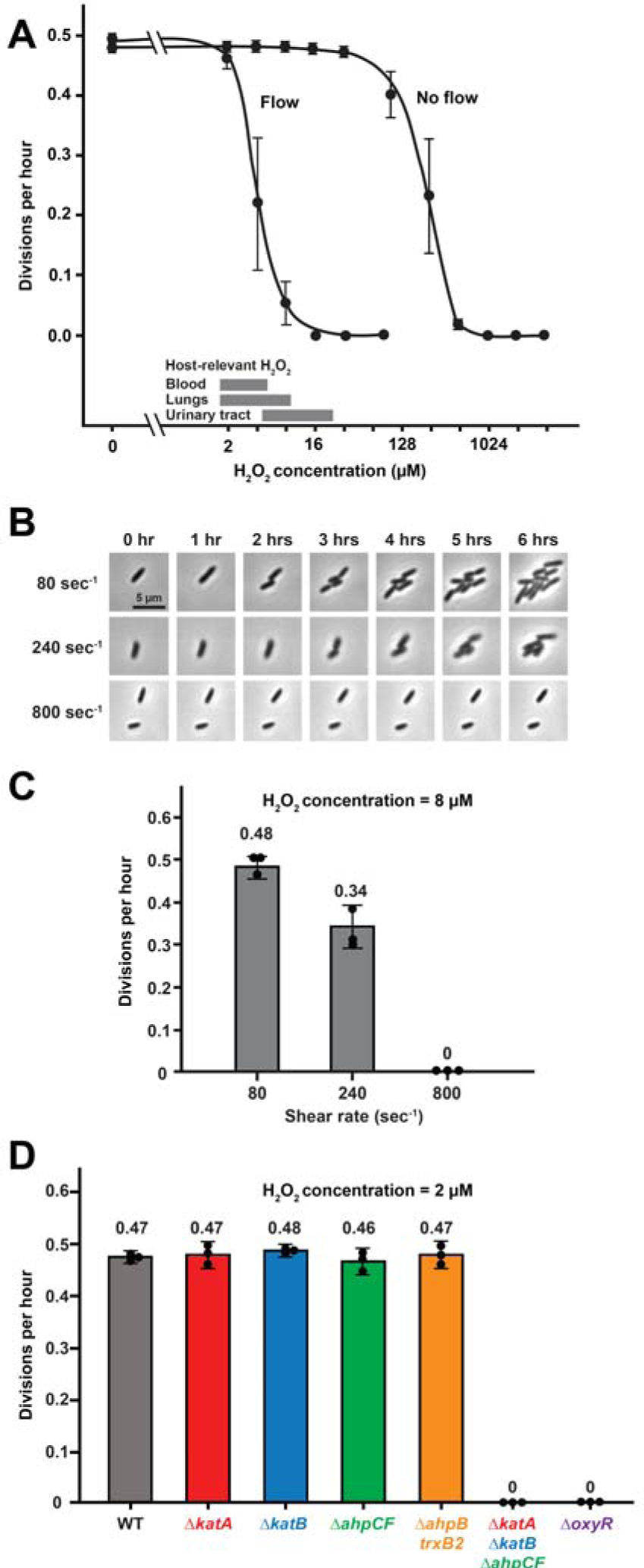
Host-relevant H_2_O_2_ and shear flow synergistically inhibit *P. aeruginosa* growth. **(A)** Quantification of wild-type cell growth over a range of H_2_O_2_ concentrations with and without flow at a shear rate of 800 sec^-1^. Gray bars represent the H_2_O_2_ concentrations found in various host tissues (21–23). **(B)** Images of cells in a microfluidic device over 6 hours with 8 µM H_2_O_2_ at shear rates of 80, 240, 800 sec^-1^. Scale bar is 5 μm. **(C)** Quantification of cell growth at different shear rates (80, 240, 800 sec^-1^) in response to 8 µM H_2_O_2._ (**D)** Quantification of cell growth in response to 2 µM H_2_O_2_ at a shear rate of 800 sec^-1^. Wild-type cells (gray), Δ*katA* (red), Δ*katB* (blue), Δ*ahpCF* (green), and Δ*ahpB*-*trxB2* (yellow) mutant cells grow in response to 2 µM H_2_O_2_. Δ*katA* Δ*katB* Δ*ahpCF* (red, blue, and green) and Δ*oxyR* (purple) mutant cells do not grow in response to 2 µM H_2_O_2_. Each biological replicate represents growth for 20 randomly chosen cells. Error bars represent SD of three biological replicates.

How does the H_2_O_2_ stress response contribute to *P. aeruginosa* growth in flow? Our model is that flow sensitizes cells to H_2_O_2_ by restoring local concentrations of H_2_O_2_ that have been depleted by cell scavenging. Thus, we hypothesize that *P. aeruginosa* H_2_O_2_ scavenging systems and the ability to upregulate these systems should be required to grow in flow. Wild-type *P. aeruginosa* cells exposed to 2 µM H_2_O_2_ with 800 sec^-1^ flow exhibited robust growth (Figure 4D). Mutant cells lacking individual scavenging systems (Δ*katA*, Δ*katB*, Δ*ahpCF* or Δ*ahpB*-*trxB2*) also exhibited robust growth, reinforcing the conclusion that these systems are redundant (Figure 4D,S5). Similar to our results with twitching motility, Δ*katA* Δ*katB* Δ*ahpCF* mutant and Δ*oxyR* mutant cells exhibited no growth (Figure 4D,S6), supporting our hypotheses that scavenging systems and their upregulation are required to grow in flow. Collectively, our experiments demonstrate that H_2_O_2_ and shear flow synergistically inhibit twitching motility and cell growth of the human pathogen *P. aeruginosa* in host-relevant conditions.

## Discussion

The ability of flow to enhance H_2_O_2_ effectiveness 50-fold reinforces the emerging idea that flow has an important impact on bacterial survival. Bacterial flow responses can be split into two categories: those that respond to mechanical forces and those that respond to chemical transport. Here, we describe a phenomenon that depends on flow-driven chemical transport. Specifically, the replenishing ability of flow overcomes cellular H_2_O_2_ removal to maintain high local concentrations and block cell growth (Figure 4). Over the course of our study, we serendipitously discovered that H_2_O_2_ replenished by flow can block *P. aeruginosa* twitching motility (Figure 3). As twitching motility is fundamentally a biological behavior dependent on mechanical forces, our work reveals a complex interplay between mechanical, chemical, and biological processes. Thus, our results demonstrate that flow has a central role in linking seemingly distinct processes and should be widely considered when designing laboratory experiments.

By combining flow and H_2_O_2_, we learned that low micromolar H_2_O_2_ levels can inhibit bacterial growth and motility (Figure 5). Our discovery counters many previous studies (38–42) that concluded micromolar H_2_O_2_ levels have no impact on bacterial growth. Based on this discrepancy, a reevaluation of experimental H_2_O_2_ concentrations is required. In previous studies, experiments were performed in static cultures, which allowed bacterial cells to protect themselves by quickly removing H_2_O_2_. In our study, flow overrode bacterial H_2_O_2_ removal, maintained constant H_2_O_2_ levels, and allowed micromolar H_2_O_2_ levels to inhibit growth. Consistent with our observations, one previous study (44) showed that manually resupplying H_2_O_2_ allowed for micromolar H_2_O_2_ to impact *E. coli*. In addition to blocking growth, our experiments reveal that micromolar H_2_O_2_ triggers a robust transcriptional response (Figure 1, Figure 2). Importantly, previous studies using much higher millimolar doses generated results that could not be fully replicated (Figure 1), suggesting that high doses generate inconsistent indirect effects. Together, these results indicate that natural micromolar H_2_O_2_ levels stress bacteria and suggest that the use of much higher millimolar H_2_O_2_ should be avoided.

**Figure 5:**
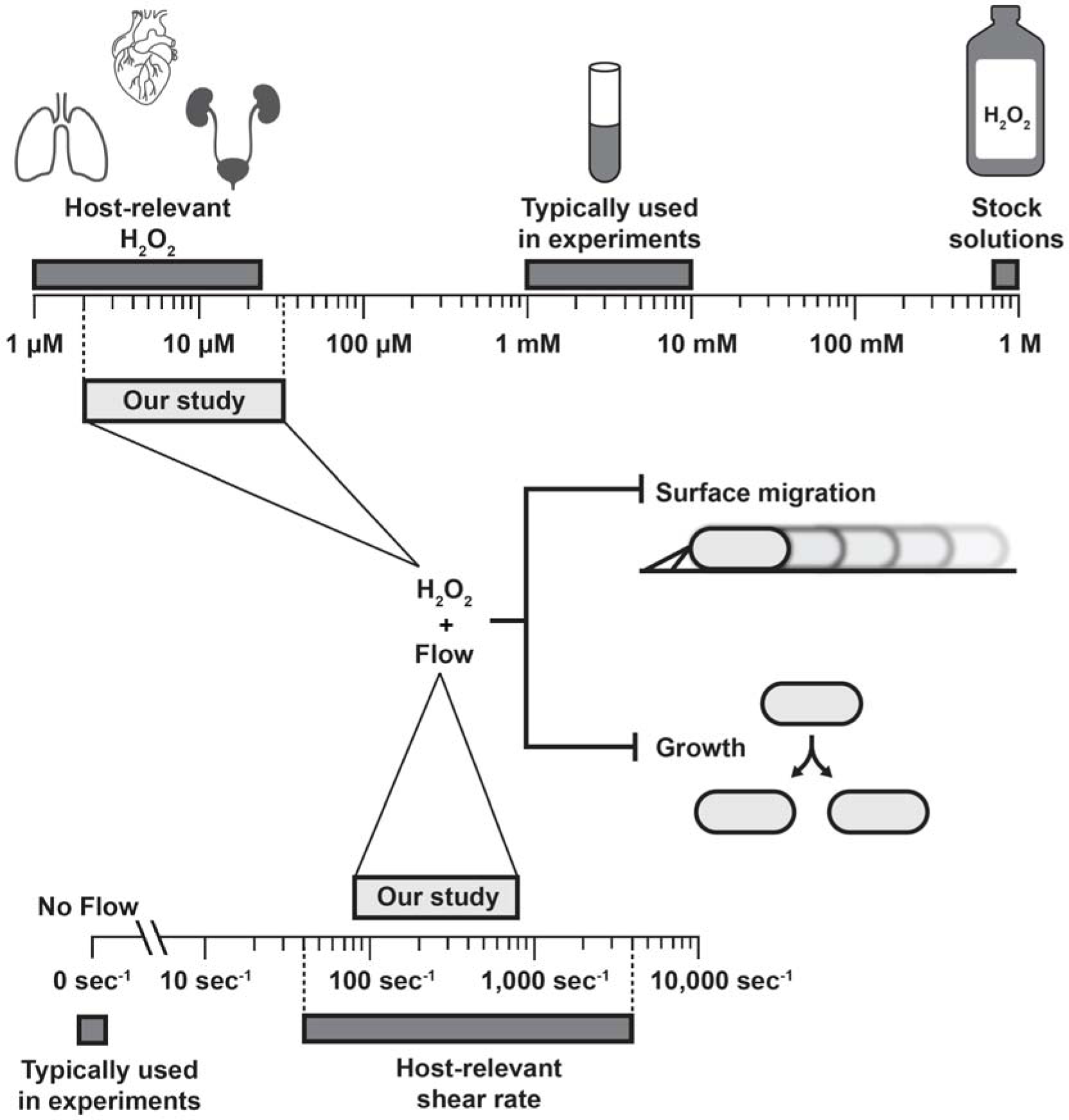
Host-relevant H_2_O_2_ and shear flow combine to inhibit migration and growth. The top scale represents H_2_O_2_ concentrations on a molar scale. It denotes concentrations found in commercially available stock solutions (0.88 M) and those typically used in laboratory experiments (1-10 mM). The left on the scale represents the reported physiological range of H_2_O_2_ in human tissues (1-25 µM)(21–23). Our study uses host-relevant H_2_O_2_ concentrations between 2-32 µM. The bottom scale represents flow in units of shear rate. Most laboratory experiments neglect flow, even though many host tissues contain flow at shear rates of 40-4,000 sec^-1^ (13–15). Our study uses host-relevant shear rates between 80-800 sec^-1^. Combining host-relevant H_2_O_2_ and flow inhibits surface migration and growth of the human pathogen *P. aeruginosa*.

Where do physical and chemical stressors co-occur in nature? Based on the ubiquity of physical and chemical stress, it is likely that they co-occur in most bacterial environments. When specifically considering flow and H_2_O_2_, the most obvious environments with both stressors are inside the human body. Specifically, the bloodstream, urinary tract, and lungs all contain flow (13–15) and micromolar H_2_O_2_ (21–23). Importantly, all of our experiments used levels of flow and H_2_O_2_ found in these environments, indicating that our experimental system is a good model of host-relevant conditions. Furthermore, *P. aeruginosa* is known to infect the bloodstream, urinary tract, and lungs, suggesting that combining flow and H_2_O_2_ has an important role in blocking bacterial growth and motility during infection. As our results provide evidence that physical and chemical stressors can have important multiplicative effects, it is now clear that we must experimentally test combinations of stressors in order to better understand natural systems.

## Acknowledgements

We thank Jessica Palalay, Piyush Sharma, and Jim Imlay for helpful discussions and comments on the manuscript. We thank Harsh Sharma for helping with RNA-sequencing analysis. We thank Alvaro Hernandez and staff at the Roy J. Carver Biotechnology Center for assistance with RNA-sequencing.

## Contributions

A.S., A.M.S., G.C.P., and J.E.S. designed research. A.S., A.M.S., and G.C.P performed research. A.S., A.M.S., G.C.P., and J.E.S. analyzed data. A.S. and J.E.S. wrote the paper.

## Supplementary Information

### Materials and Methods

#### Strains and growth conditions

The bacterial strains and plasmids used in this paper are described in Supplementary Tables S5 and S6. *P. aeruginosa* cultures were grown in either LB broth or M9 minimal medium on a cell culture roller drum, and on LB plates (1.5% Bacto Agar) at 37°C. LB broth was prepared using LB broth Miller (BD Biosciences). M9 minimal medium was prepared as 1X M9 salts, 0.4% glucose, 2 mM MgSO_4_, and 100 µM CaCl_2_.

#### Generation of *P. aeruginosa* mutants

Gene deletions were generated using the lambda Red recombinase system as previously described (6). Briefly, a deletion construct was assembled using Gibson assembly with three PCR products. First, a segment of approximately 500 bp upstream of the target insertion site was amplified from PA14 genomic DNA. Second, a fragment containing the *aacC1* ORF flanked by FRT sites was amplified from plasmid pAS03. Third, a segment of approximately 500 bp downstream of the target insertion site was amplified from PA14 genomic DNA. The three fragment Gibson product was transformed into PA14 cells expressing the plasmid pUCP18-RedS. Colonies were selected on 30 µg/mL gentamycin, followed by counter-selection of mutants of interest on 5% sucrose. Subsequently, pFLP2 was used to flip out the gentamicin antibiotic resistance gene and the colonies were selected using 300 µg/mL carbenicillin. This resulted in deletion strains with FRT sites.

#### RNA Sequencing and Analysis

RNA was isolated from mid-log cells 5 minutes after H_2_O_2_ treatment from 2 biological replicates using TRIzol^®^ Max^TM^ Bacterial RNA Isolation Kit (ThermoFisher). Preheated 200 μL Max Bacterial Enhancement Reagent was used to resuspend the cell pellet in a 1.5 mL Eppendorf tube. After incubation at 95°C for 4 minutes, 1 mL TRIzol was added followed by incubation at room temperature for 5 minutes. Phase separation was performed using 200 µL chloroform followed by centrifugation for 15 minutes at 12,000g, at 4°C. The aqueous phase ∼400 µL was then transferred to a fresh RNAse-free Eppendorf tube. The RNA was then precipitated using cold isopropanol and 75% ethanol. The RNA pellet was then air-dried before resuspending in 50 µL RNase-free water.

Genomic DNA was removed using DNase treatment. Ribosomal RNA was removed with the FastSelect Bacteria kit (Qiagen). The RNAseq libraries were prepared with the Kapa Hyper Stranded mRNAseq Sample Prep kit (Roche). Preprocessed and quality checked fastq files were obtained from the Roy J. Carver Biotechnology Center. Reads were aligned using STAR (https://github.com/alexdobin/STAR) to the publicly available *P. aeruginosa* PA14 NCBI reference genome. Aligned reads were output as BAM unsorted files, which were sorted and indexed using Samtools (https://www.htslib.org/). Gene hit counts were then calculated using featureCounts (https://subread.sourceforge.net/featureCounts.html) with gene_id as the identifier. Raw read counts of genes were normalized using trimmed mean of M-values (TMM) implemented in edgeR package (45). Differential expression was calculated using limma and the output was generated as an excel file displaying differential expression values (46).

#### Quantification of RNA levels with qRT-PCR

RNA was isolated from mid-log cells 5 minutes after H_2_O_2_ treatment as described for RNA sequencing. Quantitative real-time PCR experiments were performed using Luna® Universal One-Step RT-qPCR Kit (NEB). After thawing the components including Luna Universal One-Step Reaction Mix and primers on ice, the reaction mixture was combined by pipetting. The reaction mix was aliquoted into qPCR wells in the plate (MicroAmp™ Fast Optical 96-Well Reaction Plate, 0.1 mL, ThermoFisher). After adding the RNA template in the respective wells, the plate was sealed using MicroAmp™ Optical Adhesive Film (ThermoFisher). The plate was spun at 3,000 rpm for 1 minute to remove bubbles. The reactions were run on Applied biosystems quantitative PCR machine and all the acquired Ct values were analyzed using 5S rRNA as an internal control. Primers listed in Table S7 were used.

#### Fabrication of Microfluidic devices

Microfluidic devices were fabricated as previously described (6). Briefly, microfluidic devices were made of Polydimethylsiloxane (Dow SYLGARD 184 Kit) at a 1:10 ratio and plasma-treated to bond on a 60 mm x 35 mm x 0.16 mm superslip micro cover glass (Ted Pella, Inc.). The devices used had channels that were 500 µm wide x 50 µm tall x 2 cm long.

#### Shear rate calculations

Shear rate experienced in microfluidic devices was calculated using this equation:

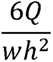

Where *Q* is flow rate, *w* is channel width, and *h* is channel height. The unit for shear rate specified as the inverse of seconds (sec^-1^).

#### Phase contrast and fluorescent microscopy

Timelapse images were captured on a Nikon ECLIPSE Ti2-E inverted microscope using the NIS Elements interface. The microscope is equipped with a Nikon 40x Plan Ph2 0.95 NA objective, and Hamamatsu ORCA-Flash4.0 LT3 Digital CMOS camera, and Lumencor SOLA Light Engine LED light source.

#### Preparation of microfluidic devices with *P. aeruginosa*

Microfluidic devices were loaded with cells as previously described (6). Briefly, all experiments were performed at approximately 22°C and with mid-log bacterial cultures. Cells were loaded into the microfluidic device using a pipette and were allowed to settle in the device for 10 minutes prior to exposure to flow. The device set-up involves the use of plastic 5 mL syringes (BD) with attached tubing connecting the needle to the inlet of the device (Brain Tree Scientific Polyethylene Tubing; ID 0.015” x OD 0.043”) that is sheathed over a 26-gauge x 1/2” hypodermic needle (Air-tite Products). These syringes were situated on a syringe pump (KD Scientific Legato 210) which was used to produce fluid flow. The outlet of the device employed the same tubing and vacated into a bleach-containing waste container. The syringe pump was used to generate flow rates of 1 - 10 µL/min, which correspond to shear rates of 80 - 800 sec^-1^.

#### Bacterial Conditioning of LB Media and H_2_O_2_ Treatment

As laboratory LB contains variable amounts of H_2_O_2_ (6), the media was conditioned as previously described (6). Briefly, *P. aeruginosa* cells were used to condition media for experiments using LB (Figure 2). Media was conditioned by diluting 50 µL of bacteria from an overnight culture into a 5 mL tube (or scaled up at the same ratio) and allowing it to sit for a defined period at 22 °C. Bacterial cells were then filtered out using a Steriflip sterile filter unit (0.22 µm pore size). To generate LB with defined H_2_O_2_ concentrations, LB was first conditioned, and then defined concentrations of H_2_O_2_ were added.

#### Quantification of *katB* expression

Expression of *katB* reporter strains was quantified similar to *fro* reporter as previously described (6). Briefly, after seeding the device with mid-log cells of *katB* reporter strain, timelapses were captured over 4 hours with pictures taken at 30-minute intervals per field (with >150 cells per field). Quantification used a MATLAB (Mathworks)-based program along with custom code to identify and quantify single cell fluorescence intensity (OUFTI). Fluorescence expression was measured of *katB*::*YFP* and constitutively expressed *P_A1/04/_*_03_-*mCherry* fluorophores. *katB* expression was then quantified as ratio of YFP to mCherry.

#### Quantification of twitching in microfluidic devices

Twitching motility was measured in M9 media with glucose in microfluidic devices in presence of flow. After seeding the devices with mid-log cells, constant M9 media with defined H_2_O_2_ concentrations was supplied to the microfluidic devices using a syringe pump. Images were taken every 5 minutes for a period of 2 hours. ImageJ software was used to visualize and track cells in the second hour of the experiment to limit any error due to initial loss of cells in flow. For each biological replicate, at least 45 cells were tracked to measure twitching. Twitching was manually defined as the displacement of a cell by at least one cell body length. Twitching motility was then quantified as the percentage of cells twitching in the entire population of cells.

#### Quantification of growth in microfluidic devices

Single-cell growth was measured in M9 media with glucose in microfluidic devices with and without flow. After seeding the devices with mid-log cells, constant M9 media with defined H_2_O_2_ concentrations was supplied to the microfluidic devices using a syringe pump. Images were taken every 5 minutes up to 6 hours. ImageJ software was used to visualize and track growth and the number of divisions were measured as the number of divisions per hour in the device. For each biological replicate, 20 cells per field at random were tracked for growth. All growth experiments were performed with cells lacking *pilA*, which prevented twitching motility and allowed for easy cell tracking.

**Table S1:** Genes upregulated after 8 μM H. _2_**O**_2_ **treatment for 5 minutes**

| Table S1: Genes upregulated after 8 μM H<sub>2</sub>O<sub>2</sub> treatment for 5 minutes | | | | |
| --- | --- | --- | --- | --- |
|  | Gene ID | Gene | P-value | log<sub>2</sub>FC |
| 1 | PA14_61040 | *katB* | 0.0002 | 6.70 |
| 2 | PA14_53300 | *ahpB* | 4.64E-06 | 6.49 |
| 3 | PA14_21530 |  | 8.98E-06 | 6.41 |
| 4 | PA14_53290 | *trxB2* | 1.62E-05 | 6.27 |
| 5 | PA14_61020 | *ankB* | 0.0043 | 5.24 |
| 6 | PA14_01720 | *ahpF* | 0.0014 | 4.17 |
| 7 | PA14_21520 |  | 0.0005 | 3.42 |
| 8 | PA14_03080 |  | 0.0017 | 3.15 |
| 9 | PA14_01710 | *ahpC* | 0.0012 | 3.07 |
| 10 | PA14_39420 |  | 0.0020 | 2.81 |
| 11 | PA14_22320 |  | 0.0087 | 2.27 |
| 12 | PA14_05870 |  | 0.0050 | 1.98 |
| 13 | PA14_38630 |  | 0.0065 | 1.98 |
| 14 | PA14_09150 | *katA* | 0.0078 | 1.96 |
| 15 | PA14_24640 |  | 0.0052 | 1.90 |
| 16 | PA14_38610 |  | 0.0034 | 1.87 |
| 17 | PA14_36770 |  | 0.0093 | 1.85 |

| Genes downregulated after 8 μM H<sub>2</sub>O<sub>2</sub> treatment for 5 minutes | | | | |
| --- | --- | --- | --- | --- |
|  | Gene ID | Gene | P-value | log<sub>2</sub>FC |
| 1 | PA14_19680 |  | 0.0007 | -2.57 |
| 2 | PA14_19690 |  | 0.0008 | -2.62 |
| 3 | PA14_64950 |  | 0.0019 | -2.69 |

**Table S2:**
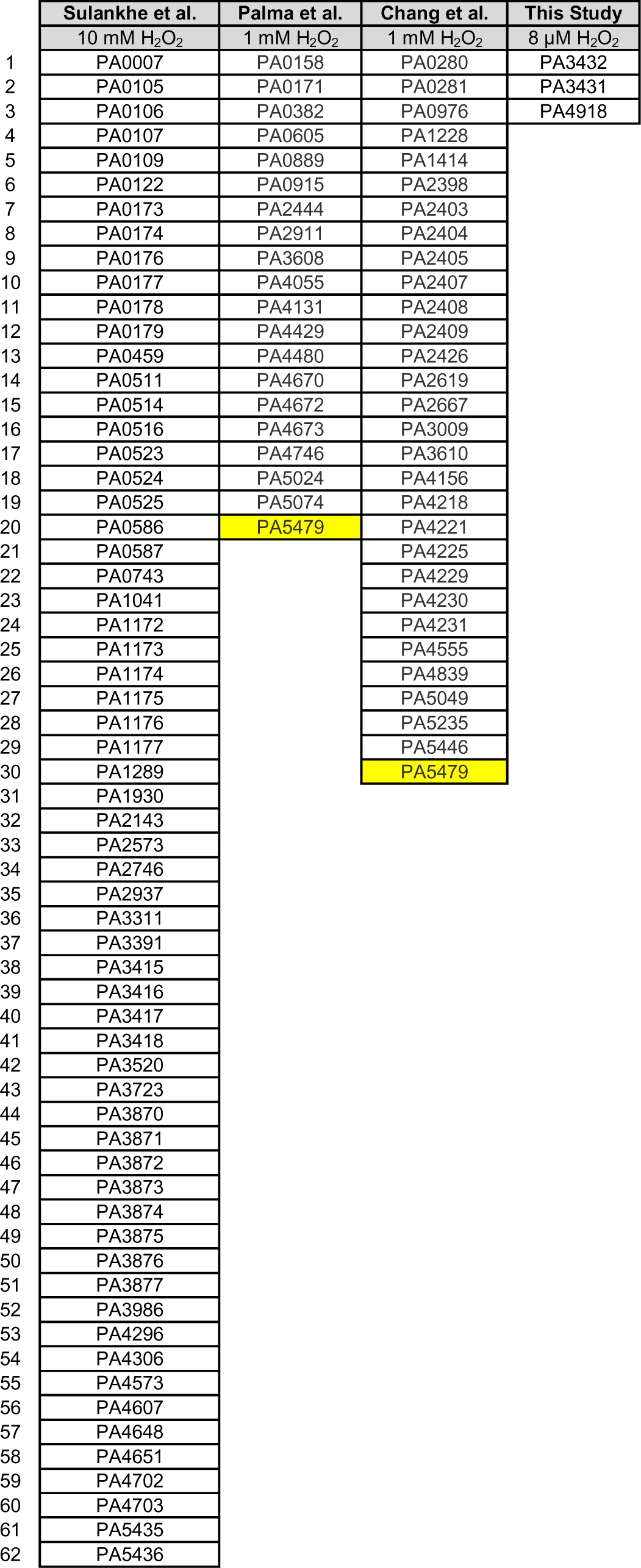
Comparative analysis of downregulated genes.

**Table S3:** Comparative analysis of upregulated genes.

| Table S3: Comparative analysis of upregulated genes |  |  |  |
| --- | --- | --- | --- |
| Sulankhe et al. | Palma et al. | Chang et al. | This Study |
| 10 mM H<sub>2</sub>O<sub>2</sub> | 1 mM H<sub>2</sub>O<sub>2</sub> | 1 mM H<sub>2</sub>O<sub>2</sub> | 8 µM H<sub>2</sub>O<sub>2</sub> |
| *katB* | *katB* | *katB* | *katB* |
| *ankB* | *ankB* | PA0182 | *ankB* |
| *ahpC* | *ahpC* | PA0612 | *ahpC* |
| *ahpF* | *ahpF* | PA0615 | *ahpF* |
| *ahpB* | *ahpB* | PA0616 | *ahpB* |
| *trxB2* | *trxB2* | PA0617 | *trxB2* |
| PA1317 | *katA* | PA0618 | *katA* |
| PA1318 | PA0250 | PA0619 | PA0249 |
| PA1320 | PA0471 | PA0620 | PA0450 |
| PA2850 | PA0472 | PA0622 | PA1942 |
| PA3237 | PA0670 | PA0625 | PA2001 |
| PA4470 | PA0671 | PA0633 | PA2002 |
|  | PA0672 | PA0634 | PA2149 |
|  | PA0962 | PA0635 | PA3050 |
|  | PA1176 | PA0636 | PA3289 |
|  | PA1230 | PA0637 | PA3237 |
|  | PA1951 | PA0638 | PA3287 |
|  | PA2025 | PA0639 |  |
|  | PA2273 | PA0641 |  |
|  | PA2426 | PA0670 |  |
|  | PA2686 | PA0671 |  |
|  | PA2687 | PA0922 |  |
|  | PA2826 | PA1466 |  |
|  | PA2827 | PA3413 |  |
|  | PA3239 | PA3414 |  |
|  | PA3600 | PA3923 |  |
|  | PA3617 | PA4582 |  |
|  | PA4219 | PA4763 |  |
|  | PA4221 | PA5470 |  |
|  | PA4227 | PA5471 |  |
|  | PA4228 | PA5530 |  |
|  | PA4230 | PA2288 |  |
|  | PA4467 | PA3007 |  |
|  | PA4468 | PA3008 |  |
|  | PA4469 |  |  |
|  | PA4471 |  |  |
|  | PA4655 |  |  |
|  | PA4764 |  |  |
|  | PA4937 |  |  |
|  | PA5532 |  |  |
|  | PA2288 |  |  |
|  | PA2850 |  |  |
|  | PA3007 |  |  |
|  | PA3008 |  |  |
|  | PA3237 |  |  |
|  | PA3287 |  |  |
|  | PA4470 |  |  |

**Table S4:** Relative transcript abundance ranking.

| Table S4: Relative transcript abundance ranking |  |  |
| --- | --- | --- |
|  | Rank out of 5,960 genes |  |
|  | No H<sub>2</sub>O<sub>2</sub> | 8 µM H<sub>2</sub>O<sub>2</sub> |
| *katA* | 73 | 10 |
| *ahpC* | 94 | 3 |
| *ahpF* | 587 | 13 |
| *katB* | 3,350 | 34 |
| *ankB* | 4,933 | 461 |
| *ahpB* | 3,978 | 143 |
| *trxB2* | 3,627 | 107 |

**Table S5:** Strains used in the study.

| Table S5: Strains used in the study |  |  |
| --- | --- | --- |
| Strain | Description | Source |
| PA14 | Wildtype; clinical isolate from burn wound | (47) |
| JS177 | $\Delta pilA::aacC1$ | (16) |
| JS176 | $\Delta pilA::FRT$ | (16) |
| JS182 | $\Delta pilTU::FRT$ | (16) |
| JS230 | $\Delta katB::aacC1$ | (6) |
| JS231 | $\Delta katB::FRT$ | (6) |
| JS232 | $\Delta ahpCF::aacC1$ | (6) |
| JS233 | $\Delta ahpCF::FRT$ | (6) |
| JS234 | $\Delta ahpCF::FRT$ pUCP18-RedS | (6) |
| JS235 | $\Delta ahpCF::FRT$ $\Delta katB::aacC1$ | (6) |
| JS236 | $\Delta ahpCF::FRT$ $\Delta katB::FRT$ | (6) |
| JS253 | $\Delta katA::FRT$ | (6) |
| JS254 | $\Delta katB::FRT$ $\Delta katA::FRT$ | (6) |
| JS255 | $\Delta ahpCF::FRT$ $\Delta katA::FRT$ | (6) |
| JS256 | $\Delta ahpCF::FRT$ $\Delta katB::FRT$ $\Delta katA::FRT$ | (6) |
| JS270 | $P_{katB}::[YFP-FRT]$ $attB::[plac-mcherry FRT]$ | This paper |
| JS277 | $P_{katB}::[YFP FRT]$ $attB::[plac-mcherry FRT]$ $\Delta oxyR::aacC1$ | This paper |
| JS240 | $\Delta oxyR::aacC1$ | This paper |
| JS241 | $\Delta oxyR::FRT$ | This paper |
| JS285 | $\Delta oxyR::FRT$ pUC18RedS | This paper |
| JS296 | $\Delta ahpBtrxB2::aacC1$ | This paper |
| JS320 | $\Delta ahpBtrxB2::FRT$ | This paper |
| JS287 | $\Delta oxyR::FRT$ $\Delta pilA::aacC1$ | This paper |
| JS323 | $\Delta katA::FRT$ $\Delta pilA::aacC1$ | This paper |
| JS324 | $\Delta katA::FRT$ $\Delta pilA::FRT$ | This paper |
| JS290 | $\Delta katB::FRT$ $\Delta pilA::aacC1$ | This paper |
| JS291 | $\Delta ahpCF::FRT$ $\Delta pilA::aacC1$ | This paper |
| JS279 | $\Delta ahpBtrxB2::FRT$ $\Delta pilA::aacC1$ | This paper |
| JS288 | $\Delta ahpCF::FRT$ $\Delta katB::FRT$ $\Delta katA::FRT$ $\Delta pilA::aacC1$ | This paper |

**Table S6:** Plasmids used in the study.

| Table S6: Plasmids used in the study |  |  |
| --- | --- | --- |
| Plasmid | Description | Source |
| pAS03D | Plasmid for generating deletion mutants in *P. aeruginosa* | (48) |
| pFLP2 | Plasmid expressing FLP2 to recombine FRT sites | (49) |
| pUCP18-RedS | Lambda red recombineering vector | (48) |

**Table S7:**
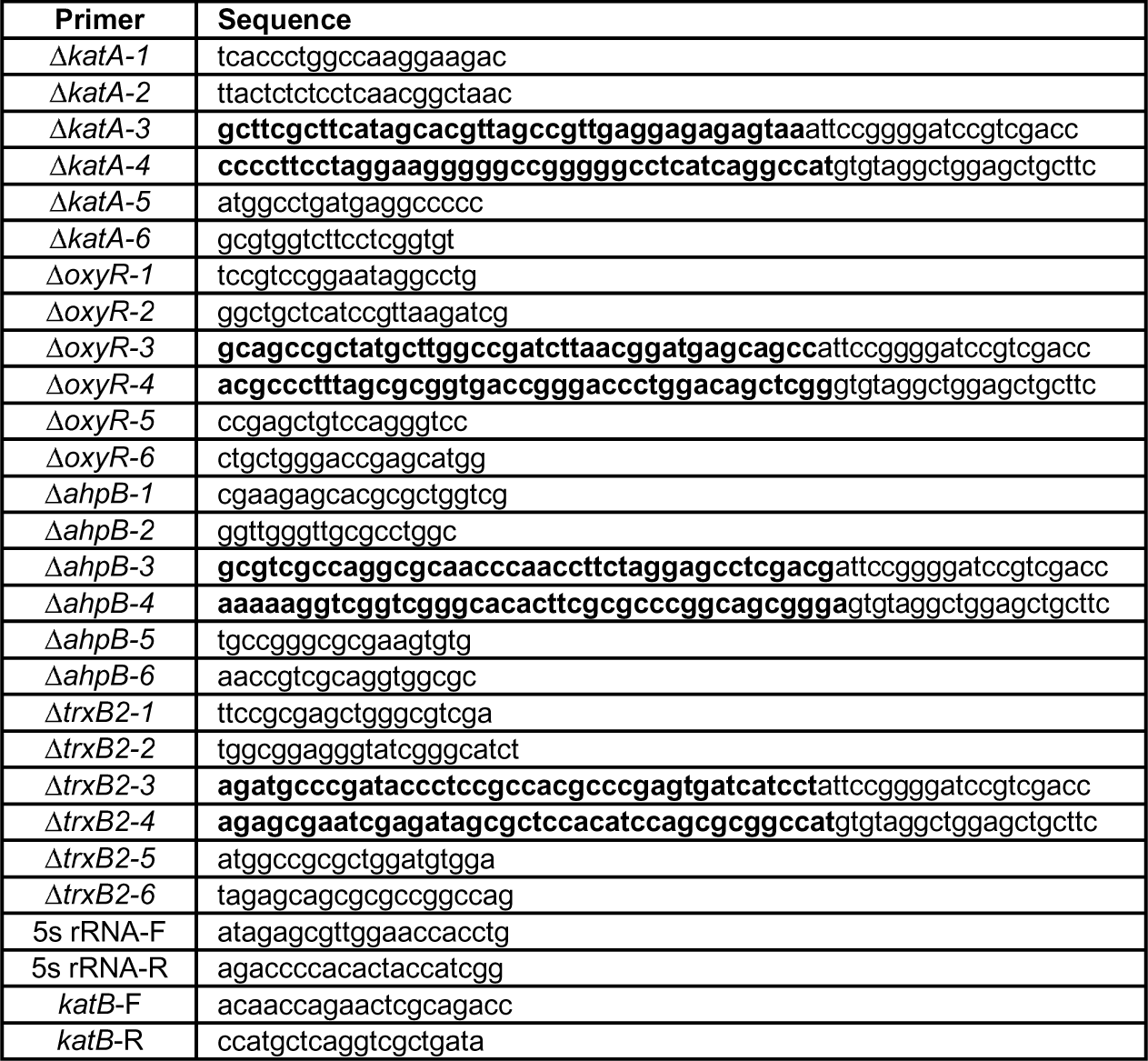
Primers used in this study.

**Figure S1:**
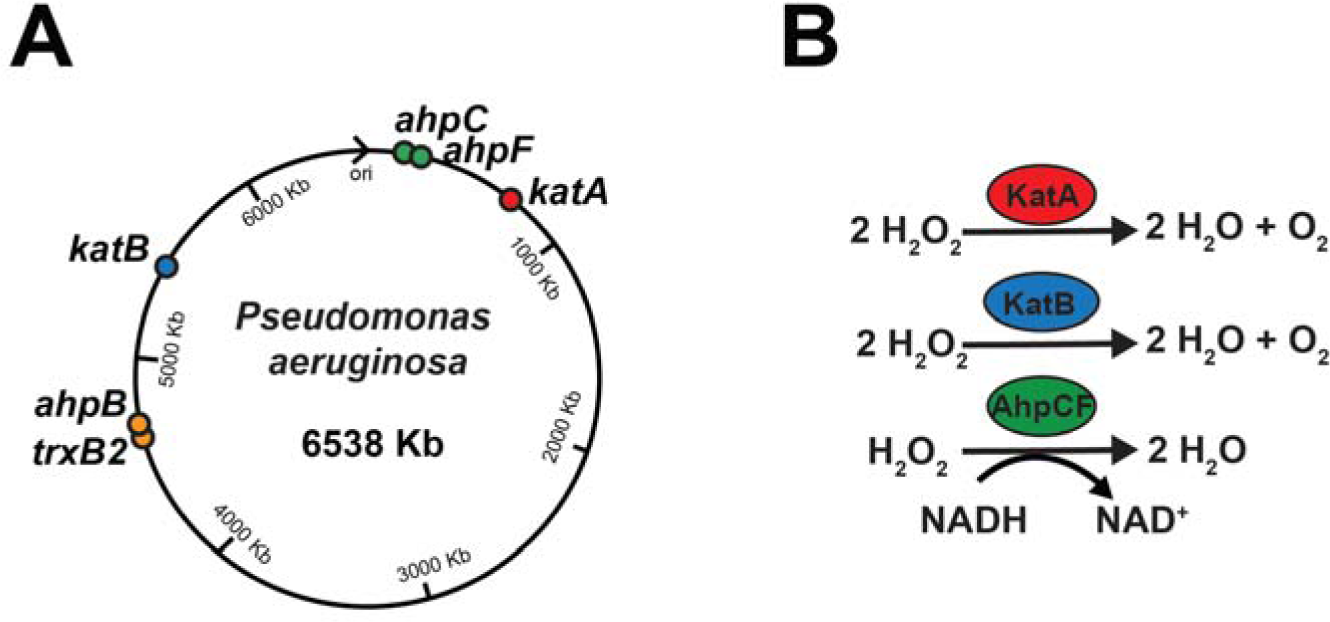
Genome localization and function of *P. aeruginosa* scavenging systems. **(A)** Localization of H_2_O_2_-sensitive genes on the *P. aeruginosa* PA14 genome. H_2_O_2_-sensitive genes cluster into 4 distinct genomic locations. *katA* and *ahpCF* are located close to the origin of replication (ori), consistent with their very high expression levels. **(B)** Biochemical reactions carried out by H_2_O_2_ scavenging enzymes. KatA, KatB, and AhpCF scavenge H_2_O_2_. KatA and KatB are catalases and break down H_2_O_2_ into water and O_2_. AhpC and AhpF works as a pair to reduce H_2_O_2_ to water using NADH as a reductant.

**Figure S2:**
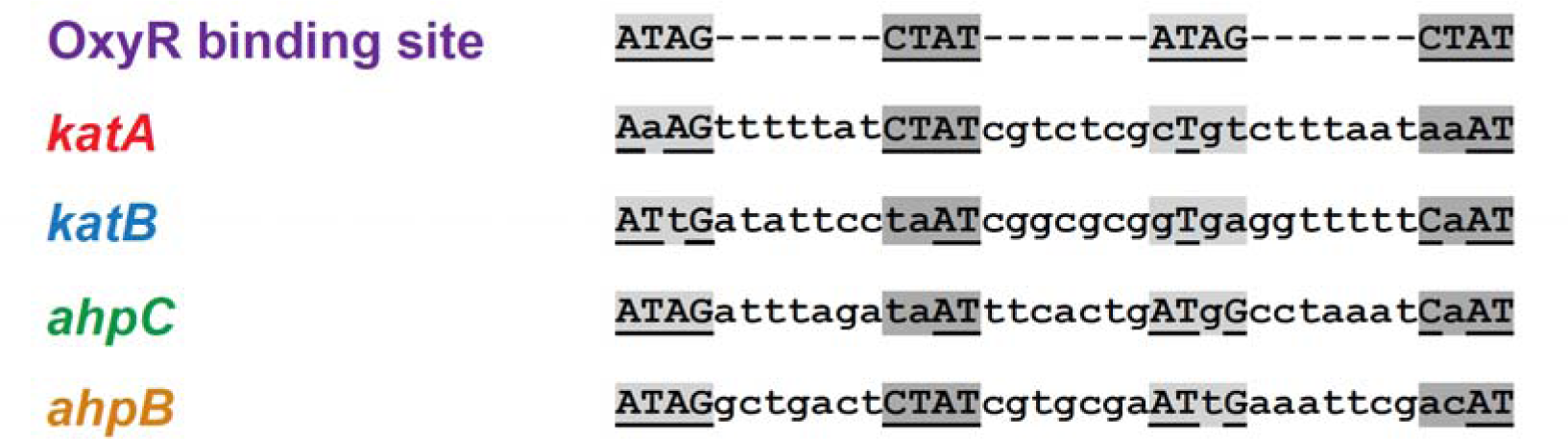
Regions upstream of scavenging genes contain OxyR binding sites. Upstream sequences of H_2_O_2_-sensitive genes contain binding sites for OxyR. The OxyR binding site is reported to be four tetra-nucleotide sequences, each separated by 7 nucleotides (26, 33). The alignment of the OxyR-regulated promoters highlights four OxyR-binding sequences (highlighted in gray). Residues that align with the consensus sequence of OxyR binding site are underlined. The number of bases matching the OxyR binding sequence are 10 of 16 (*katA*), 9 of 16 (*katB*), 12 of 16 (*ahpC*), and 13 of 16 (*ahpB*).

**Figure S3:**
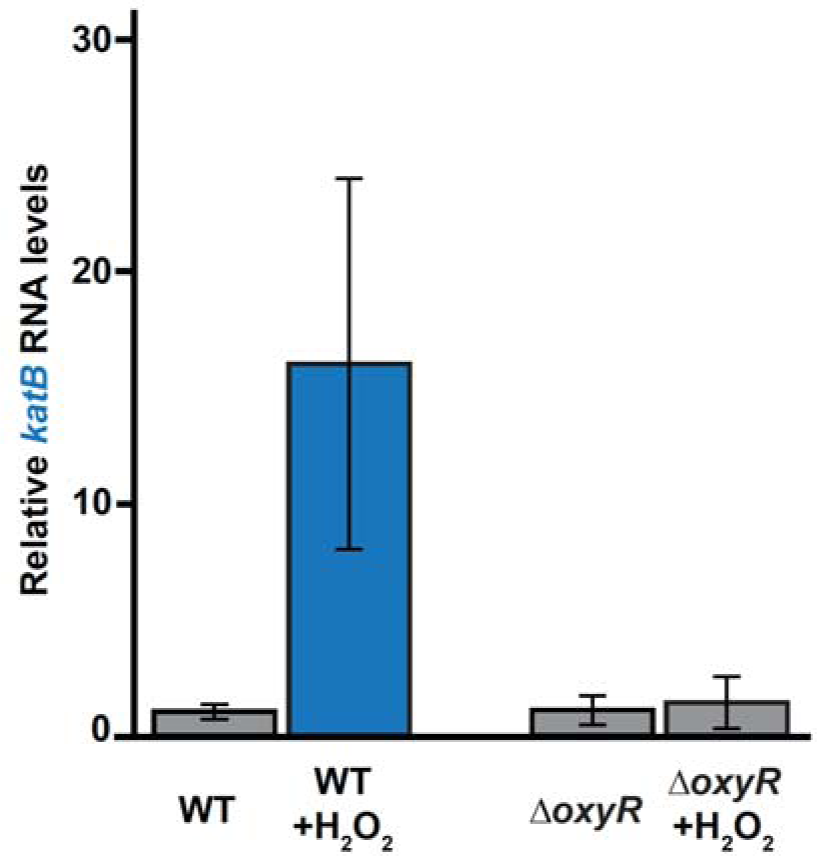
OxyR is required for H_2_O_2_ induction of *katB* RNA levels. *katB* RNA levels in wild-type and Δ*oxyR* cells from a qRT-PCR experiment with and without 8 µM H_2_O_2_. In the presence of H_2_O_2_, wild-type cells show induction of *katB* RNA levels in response to H_2_O_2_. In contrast, Δ*oxyR* mutant cells exhibit no induction of *katB* RNA levels in response to H_2_O_2._ These results confirm that *katB* levels are regulated by H_2_O_2_ and OxyR. Error bars represent SD of three biological replicates.

**Figure S4:**
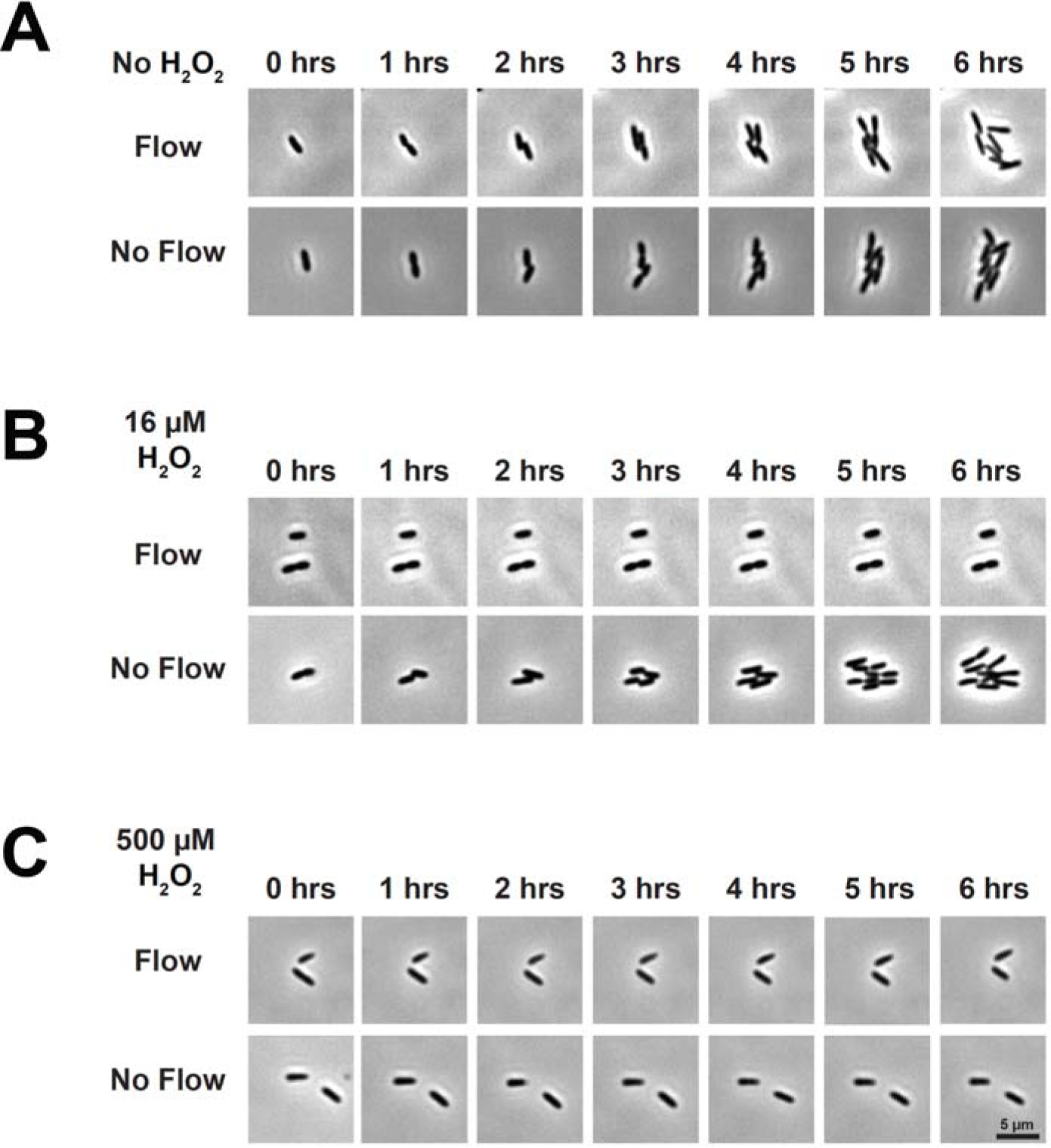
Host-relevant H_2_O_2_ and flow synergistically inhibit *P. aeruginosa* growth. **(A)** Images of cells in a microfluidic device over 6 hours without H_2_O_2_ at shear rates of 0 and 800 sec^-1^. Cells grow with and without flow. **(B)** Images of cells in a microfluidic device over 6 hours with 16 µM H_2_O_2_ at shear rates of 0 and 800 sec^-1^. Cells do not grow in flow but grow without flow. **(C)** Images of cells in a microfluidic device over 6 hours with 500 µM H_2_O_2_ at shear rates of 0 and 800 sec^-1^. Cells do not grow with and without flow. Images are representative examples of quantification of cell growth in Figure 4A. Scale bar is 5 μm.

**Figure S5.**
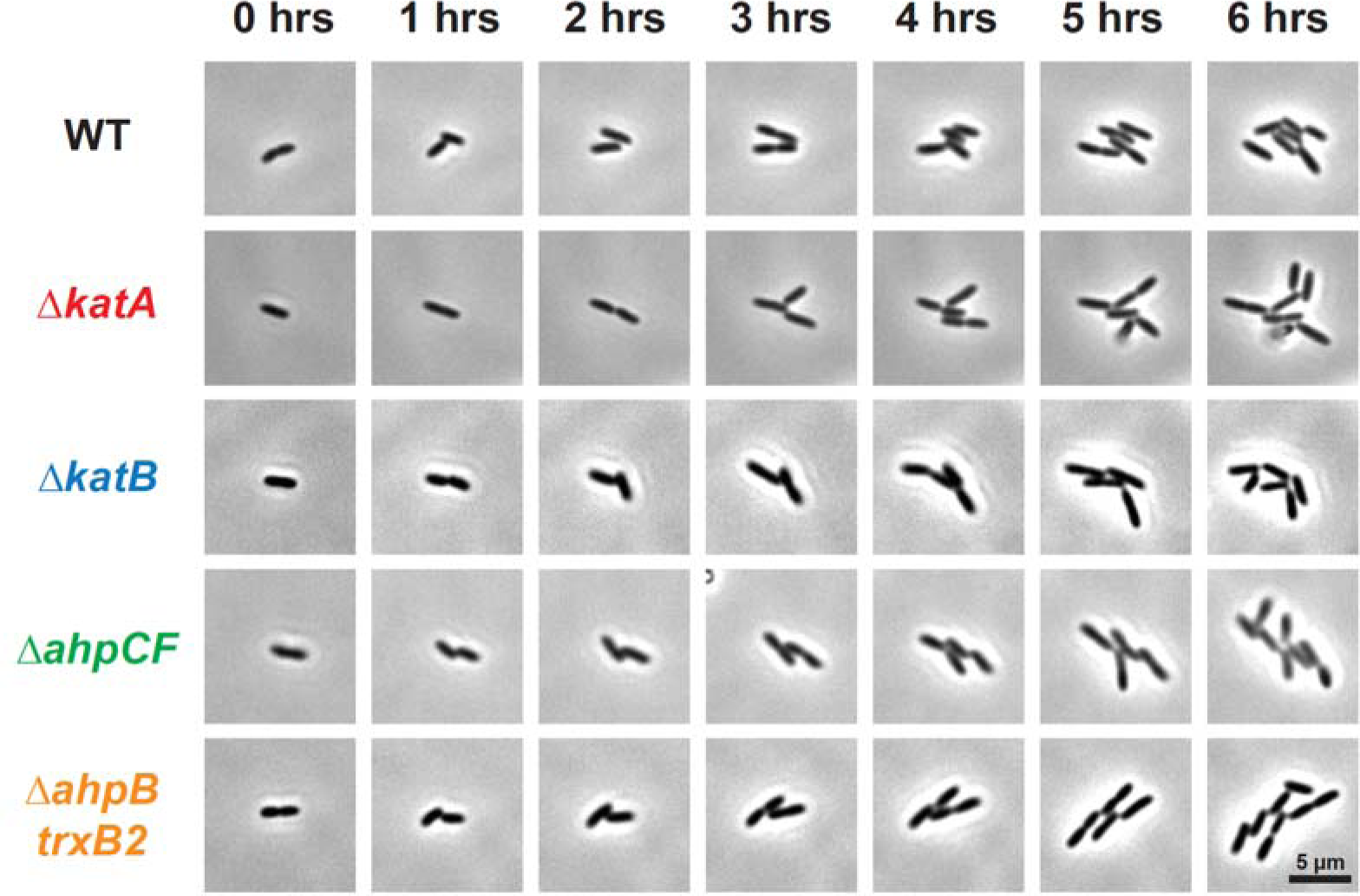
*P. aeruginosa* scavenging systems have redundant roles. Images of cells in a microfluidic device over 6 hours with 2 µM H_2_O_2_ at shear rate of 800 sec^-1^. Cells lacking single scavenging systems exhibit indistinguishable growth from wild-type cells. Images are representative examples of quantification of cell growth in Figure 4D. Scale bar is 5 μm.

**Figure S6:**
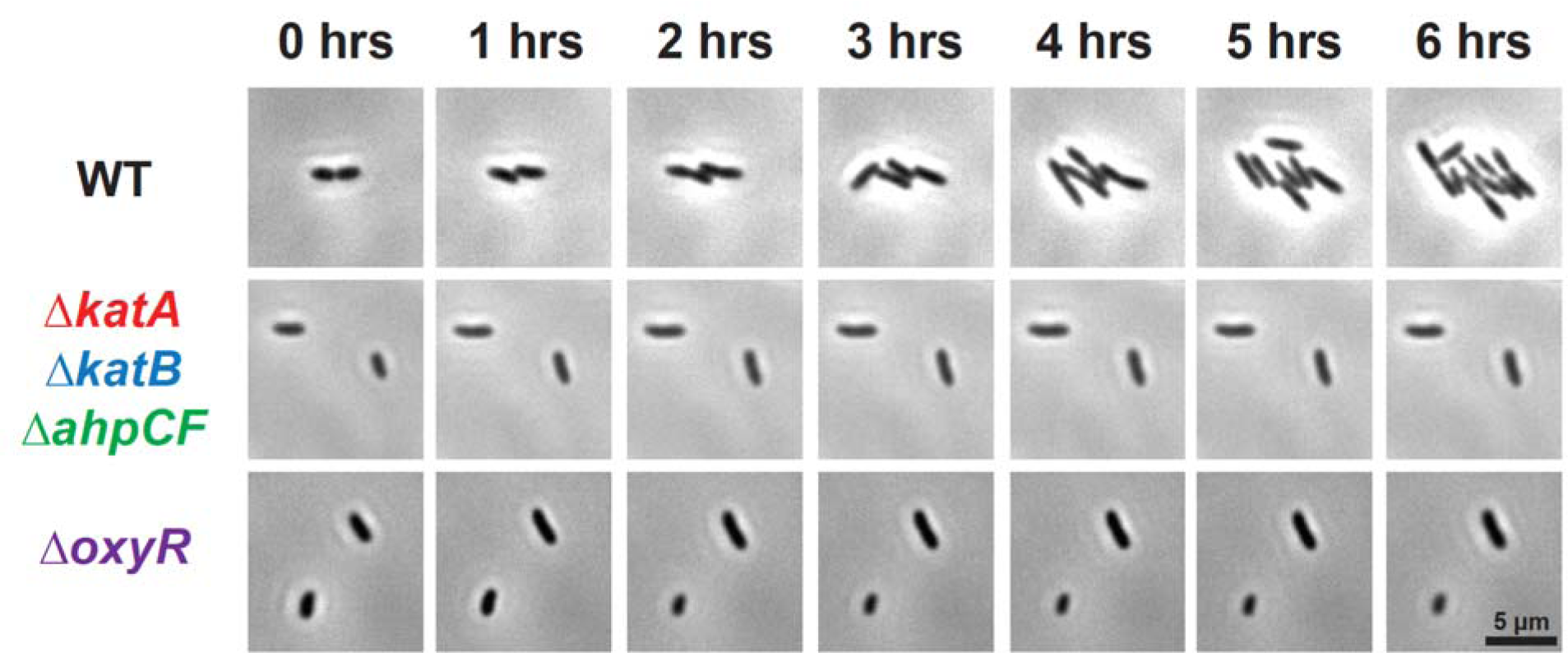
*P. aeruginosa* scavenging systems and OxyR are required to grow in conditions with H_2_O_2_ and flow. Images of cells in a microfluidic device over 6 hours with 2 µM H_2_O_2_ at shear rates of 800 sec^-1^. Wild-type cells grow while Δ*katA* Δ*katB* Δ*ahpCF* and Δ*oxyR* mutant cells do not grow in response to 2 µM H_2_O_2_. Images are representative examples of quantification of cell growth in Figure 4D. Scale bar is 5 μm.

